# Multiple exposures to sevoflurane during the neonatal period impair hippocampus development and cognitive function in young mice

**DOI:** 10.1101/2023.07.02.547420

**Authors:** Gaojie Yu, Xiaoting Huang, Liang Lin, Zhenyi Chen, Jie Zhang, Jia Jia

## Abstract

Sevoflurane is a commonly used anesthesia for infants and young children. Sevoflurane has potential neurotoxicity in immature brains; however, its specific mechanism has not been fully elucidated. Therefore, we used an established sevoflurane anesthesia model and evaluated hippocampal synaptic function using transcriptome sequencing, biochemical analyses, and animal behavior to investigate the effect of multiple neonatal sevoflurane exposures on the hippocampus in mice. C57BL/6J mice were randomly divided into sevoflurane and control groups. All experimental conditions were identical in the two groups except for the anesthetization procedure. Mice in the sevoflurane group were anesthetized with 2.5% sevoflurane for 2 h daily for 3 consecutive days (postnatal days 6−8). Mice in the control group did not receive sevoflurane anesthesia. During anesthesia, mice were administered 50% oxygen, and the respiratory rate and skin color were monitored. On day 3 of modeling, half of the mice were randomly selected to undergo harvesting of the hippocampus. RNA sequencing (RNA-seq) of RNA extracted from the hippocampus identified 736 differentially expressed genes (DEGs), including 433 upregulated and 303 downregulated DEGs, after multiple sevoflurane exposures. Gene ontology term enrichment analysis results suggested that sevoflurane exposure altered the expression of neurodevelopment-related genes in neonatal mice. Several enriched biological processes involved in brain development (axon/forebrain development) and adenosine monophosphate-activated protein kinase signaling pathways were highlighted. Comparison with RNA-seq database information showed that DEGs of the neonatal hippocampus after multiple exposures to sevoflurane were specific to neonatal mice. Furthermore, Morris water maze testing confirmed that sevoflurane anesthesia induced learning and memory impairments in young mice. Additionally, Western blot and immunofluorescence analyses showed that sevoflurane treatment decreased synaptic protein levels, such as postsynaptic density protein 95, synaptosomal-associated protein, 25 kDa, and B-cell lymphoma 2-associated athanogene 3, in the hippocampus, which induced synaptic dysfunction, resulting in impaired nervous system development in young mice.

## Introduction

There is a growing need for infants and young children to undergo general anesthesia for surgery or clinical examination. However, the effects of general anesthetic agents on the function and structure of the brain during early-stage development are unclear. The New York Medical Database suggests that general anesthesia may be toxic to the neurodevelopment of infants 0−3 years of age [1]. The United States Food and Drug Administration issued a drug safety statement concerning the use of 11 commonly prescribed sedative and anesthetic drugs, including sevoflurane, with potential neurotoxic effects on children younger than 3 years of age [2]. A previous study also indicated that clinicians should carefully formulate anesthesia plans for children who must receive anesthesia [3]. A study by Mayo Anesthesia Safety in Kids indicated that slight declines in fine-motor coordination and processing speed might impede learning after sevoflurane exposure [4]. The neurotoxicity and side effects of general anesthesia have received increasing attention; therefore, it is necessary to research and elucidate any potential negative effects on the neurodevelopment of children.

Sevoflurane is neurotoxic to the developing brain when used repeatedly or for long periods. Chen et al. found that the offspring of female rats exposed to anesthesia with sevoflurane during pregnancy had social interaction deficits [5]. In mice, sevoflurane may increase the risk of cognitive dysfunction and induce impulsive behavior in adulthood, similar to that associated with attention-deficit/hyperactivity disorder [6]. Various studies have demonstrated that multiple exposures to sevoflurane in neonatal mice significantly altered the genome-wide expression profile of different brain regions in young mice [7], rats [8], and monkeys [9]. However, these studies only investigated the expression of differentially expressed genes (DEGs) in young animals. No study has identified neonatal hippocampal transcriptome alterations through high-throughput RNA sequencing (RNA-seq).

Therefore, we investigated the mechanism of sevoflurane-induced neurotoxicity in the newborn hippocampus using a well-established model of multiple exposures to sevoflurane during the neonatal period. Furthermore, we evaluated the influence of sevoflurane on cognitive function behaviors in young mice. The main objective of this study was to demonstrate the genome-wide expression profile and DEGs of the neonatal hippocampus of sevoflurane-treated and control mice using RNA-seq. Therefore, enrichment of biological processes and signaling pathways of DEGs were analyzed. Western blotting and immunostaining were performed to further elucidate the molecular mechanism by which sevoflurane induces abnormal brain development and cognitive dysfunction.

## Materials and methods

### Overview

This study was designed to investigate the effects of sevoflurane on the hippocampus of neonatal mice and brain function (Fig 1A). Neonatal mice were exposed to sevoflurane for 2 h on 3 consecutive postnatal days (postnatal days 6–8). On postnatal day 8, the hippocampus was extracted for transcriptome analysis using RNA-seq, and synaptic proteins were analyzed using Western blot and immunostaining. The Morris water maze test was performed on postnatal days 38−44 to assess cognitive function behaviors in young mice after neonatal sevoflurane exposure. To implement an unbiased analysis of the effects of multiple exposures to sevoflurane on the neonatal hippocampus, we conducted gene expression profiling in the neonatal hippocampus by performing a genome-wide transcription analysis with next-generation sequencing and RNA-seq analysis.

**Fig 1.**
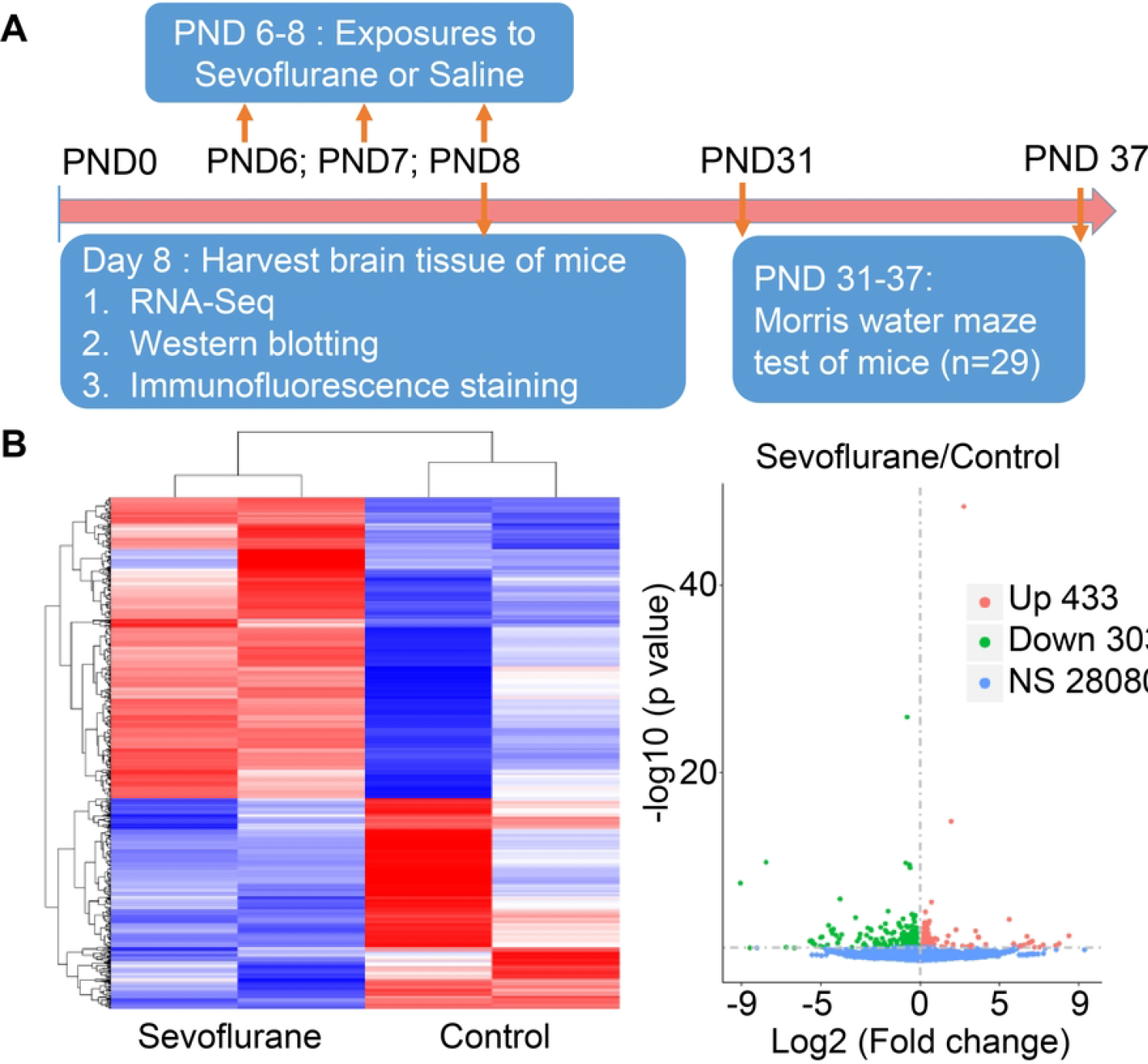
Study schema, heatmap, and volcano plot. A) Schematic model of this study. Exposure to sevoflurane for 2 h on 3 consecutive days (postnatal days [PNDs] 6−8). Samples were harvested for RNA sequencing, Western blotting, and immunostaining. The Morris water maze was performed on PND 31. B) Heatmap of upregulated and downregulated genes. C) Volcano plot illustrating upregulated (red dots) and downregulated (green dots) genes and normally expressed genes (blue dots).

### *In vivo* experiments

C57BL/6J mice were used in this study. The breeding and experimental procedures complied with the regulations of the Animal Ethics Committee of Xiamen University. A 12-h light/12-h dark cycle was applied in the room where the mice were housed. The room temperature ranged from 22°C to 25°C, with 50% humidity. On days 6−8 after birth (postnatal days 6−8), the mice were selected for further analyses.

### Establishment of the anesthesia model

A gas analyzer was used to monitor the concentrations of sevoflurane and oxygen throughout the experiment, and a heating pad was placed under the anesthesia room to maintain a rectal temperature of 37.5°C (±0.5°C) in the mice. Mice were randomly divided into sevoflurane and control groups. Mice in the sevoflurane group were exposed to 2.5% sevoflurane for 3 consecutive days (postnatal days 6−8). The flow rate was 2 L/min for the first 5 min of anesthesia induction; thereafter, it was maintained at 1 L/min. During anesthesia, 50% oxygen was administered, and the respiratory rate and skin color of the mice were monitored. The control group was not administered sevoflurane anesthesia; however, the other experimental conditions were the same. On day 3 of sevoflurane anesthesia, after the mice woke, half of them were randomly selected to undergo harvesting of the hippocampus, which was stored in a -80°C refrigerator. The remaining mice were kept for animal behavioral experiments.

### Morris water maze

Researchers who performed the Morris water maze test touched the mice on 3 consecutive days. This allowed the mice to adapt to the researchers. Experiments were conducted on postnatal days 38−44. The mice were kept in a quiet environment to avoid shock.

There were 15 and 14 mice in the control and sevoflurane groups, respectively. All mice grew normally, and there was no significant difference in body weight between groups. The circular pool of the experiment was divided into four equal 90° quadrants (southwest, southeast, northeast, and northwest) using software. A platform was placed 1 cm under the water in the central area of the southwest quadrant; this area was the target zone. During postnatal days 38−43, mice were trained to familiarize themselves with the pool environment. Specifically, two water entry points were randomly selected from the four quadrants to begin training. If the mice did not reach the target platform within 1 min, they were guided manually and kept on the platform for an additional 10 s. Escape latency was calculated as the time from when the mice entered the water to the time when they reached the platform.

The mice were evacuated from the underwater platform on postnatal day 44 for the space exploration ability test. The starting point of each mouse was in the opposite direction of the platform. The exploration trajectory of each mouse in the pool was observed and recorded, including the time required to cross the platform for the first time, the number of times the platform area was crossed, and the time spent in the target quadrant.

### Western blot experiment

Total proteins were extracted and separated by 10% sodium dodecyl sulfate-polyacrylamide gel electrophoresis. They were then electrophoretically transferred to a polyvinylidene fluoride membrane. After blocking in 5% nonfat milk for 1 h at room temperature (20–25°C), the membranes were incubated with appropriately diluted primary antibodies overnight at 4°C. After washing with Tris-buffered saline containing Tween 20, the membranes were incubated with horseradish peroxidase-conjugated secondary antibody (1:20,000 dilution) for 1 h at 20−25°C. The membranes were visualized using an enhanced Phototope TM-HRP Detection Kit and exposed to a Bio-Rad processor. The antibodies used during this study were β-actin (1:1000; Abcam, Cambridge, UK), PSD95 (1:1000; Abcam), Bag3 (1:1000; Abcam), SNAP25 (1:1000; Abcam), and p-Bag3 (1:1000; Abcam).

### Tissue immunofluorescence

Mouse brains were placed in 4% paraformaldehyde and frozen at 4°C for 24 h.Sucrose solutions (20%, 25%, and 30%) were sequentially replaced every 24 h for gradient dehydration. After the cerebrum was removed, it was embedded in an optimal cutting temperature compound, and the slices of the classic hippocampal structure were obtained using a freezing microtome. The slides were removed, rinsed four times with 1× phosphate-buffered saline, and permeabilized with 0.3% TritonX-100 for 15 min. Antigen retrieval was performed with 1× citric acid retrieval solution, and cells were blocked at room temperature for 1 h. Bag3 antibody (1:1000; Abcam, UK) was incubated at 4°C overnight, and the secondary antibody was incubated at room temperature for 1 h. Thereafter, 4’,6-diamidino-2-phenylindole stain was applied for 1 min, and photography was performed under a fluorescence microscope.

### Statistical analysis

Image J (National Institutes of Health, Bethesda, MD), Photoshop 6.0 (Adobe Systems, San Jose, CA), GraphPad Prism 9 (GraphPad Software, San Diego, CA), and other statistical software were used for processing, quantitative analyses, and data graphing. The t-test and variance analysis were used to evaluate the significance of data differences. P<0.05 was considered statistically significant.

## Results

### Quality control of RNA-seq data and gene alignment analysis

After comparing the clean reads, 74,969,859 and 65,510,332 raw reads were obtained for control 1 and control 2 subgroups, with rates of 95.85% and 95.09%, respectively.

Regarding the sevoflurane group, 79,645,621 and 59,076,326 total useful reads were obtained for sevoflurane 1 and sevoflurane 2 subgroups, with match rates of 95.85% and 96.56%, respectively.

### Transcriptome analysis of differential gene expression

RNA-seq analysis showed the expression level of 28,816 genes in the neonatal hippocampus. Among these, the expression levels of 433 genes were significantly increased in mice exposed to sevoflurane, whereas those of 303 genes were significantly decreased (Fig 1B, C). A heat map displaying the transcript clusters with differential expression in the neonatal hippocampus emphasized that these DEGs distinguished the sevoflurane group from the control group, suggesting that gene expression in the hippocampus was dramatically influenced by multiple exposures to sevoflurane.

### Gene ontology analysis of DEGs

Based on the DEGs of the sevoflurane and control groups, we performed a functional analysis with three main ontologies: biological processes, molecular functions, and cellular components. The 10 main biological processes, molecular functions, and cellular components after functional and enrichment analyses are shown in Fig 2. Regarding the biological processes of upregulated DEGs, those involved in brain development included axon, telencephalon, forebrain, and pallium development. Those associated with downregulated DEGs included extracellular matrix/structure organization, potassium ion transport, and oligodendrocyte differentiation.

**Fig 2.**
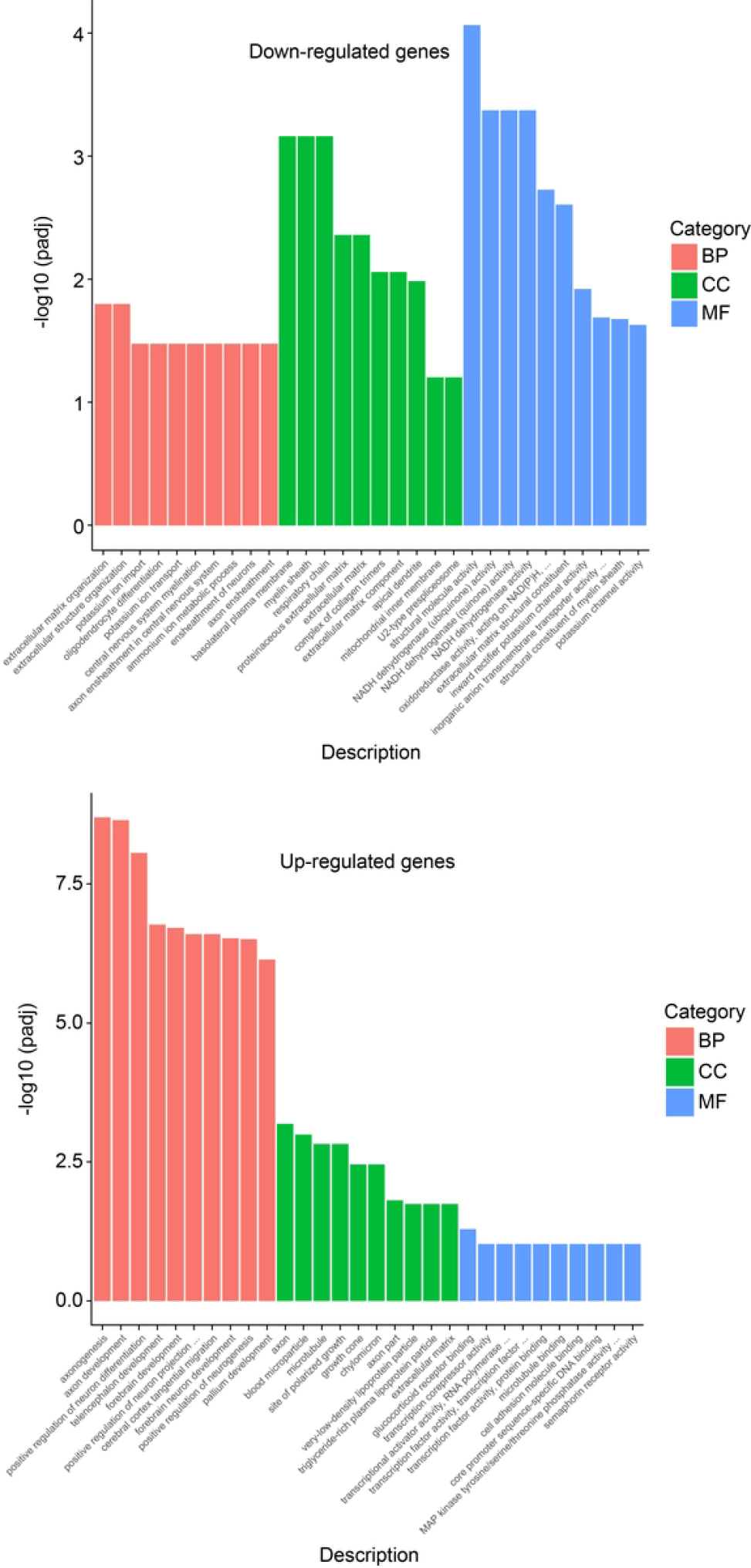
Gene ontology analysis of differentially expressed genes between the sevoflurane and control groups. The 30 most significant pathways from gene ontology (GO) enrichment analysis were selected for this bar graph. The abscissas represent the genetic pathways in GO analysis, and the ordinates represent the significance level of pathway enrichment. Red represents biological processes (BPs), green represents cellular components (CCs), and blue represents molecular function (MF). The top graph contains the 30 most downregulated pathways, and the bottom graph contains the 30 most upregulated pathways.

The enrichment of DEG Gene Ontology (GO) terms was also classified based on the cellular components and molecular functions. Regarding cellular components, the most significant categories were basolateral plasma membrane for downregulated DEGs and axons for upregulated DEGs. Regarding molecular functions, the most significant categories were glucocorticoid receptor binding for upregulated genes and structural molecule activity for downregulated genes.

### DEG pathway analysis revealed significantly enriched signaling pathways

We performed a Kyoto Encyclopedia of Genes and Genomes (KEGG) pathway analysis of the DEGs. Fig 3 shows the most enriched pathways for upregulated and downregulated genes. Retrograde endocannabinoid signaling, oxidative phosphorylation, and Parkinson’s disease were the three most enriched pathways associated with downregulated DEGs. The three most enriched pathways associated with upregulated DEGs were the adenosine monophosphate-activated protein kinase (AMPK) signaling pathway and signaling pathways regulating the pluripotency of parathyroid hormone synthesis.

**Fig 3.**
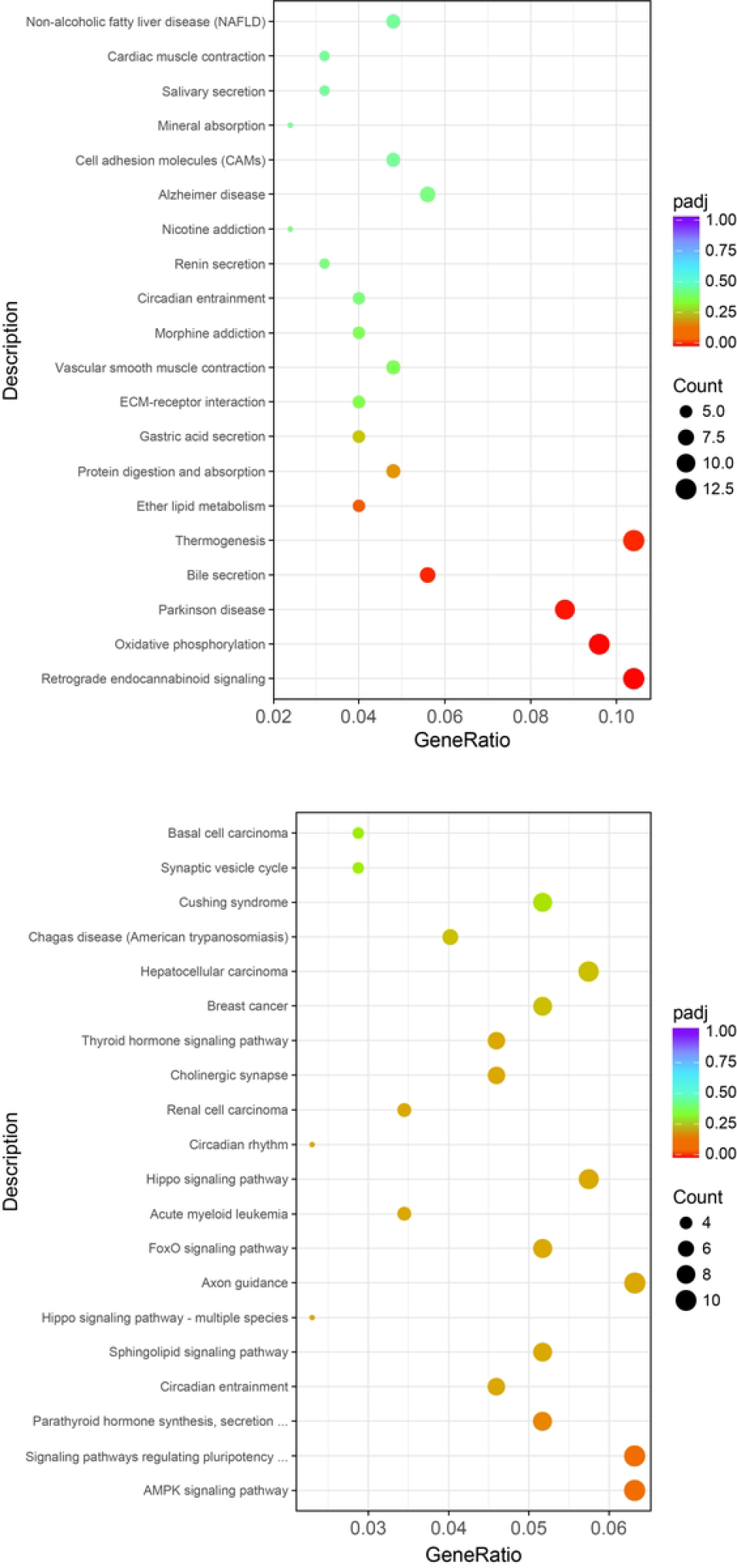
KEGG pathway analysis of differentially expressed genes. The top panel illustrates the most enriched pathways for downregulated genes. The bottom panel illustrates the most enriched pathways for upregulated genes.

### Comparison of the transcriptome DEG dataset after sevoflurane exposure

To identify specific gene expression changes caused by sevoflurane exposure in the neonatal period, transcriptome array data of the hippocampus of young mice treated with sevoflurane were compared with data from two studies by Yamamoto et al. (referred to as Young1) [10] and Song et al. (referred to as Young2) [7]. We chose the young mouse hippocampal data from Young1 because our data focused on the hippocampus. Comparison of the DEGs of our neonatal mice and the two published datasets on the young mouse hippocampus (Fig 4) revealed only nine and five downregulated DEGs in common with Young 1 and Young2 data, respectively. In addition, there were six and three upregulated DEGs in common with Young1 and Young2 data, respectively. This comparison of the upregulated and downregulated DEGs showed that the neonatal hippocampal DEGs, with the exception of several common genes, were expressed differently in the young hippocampus after sevoflurane treatment.

**Fig 4.**
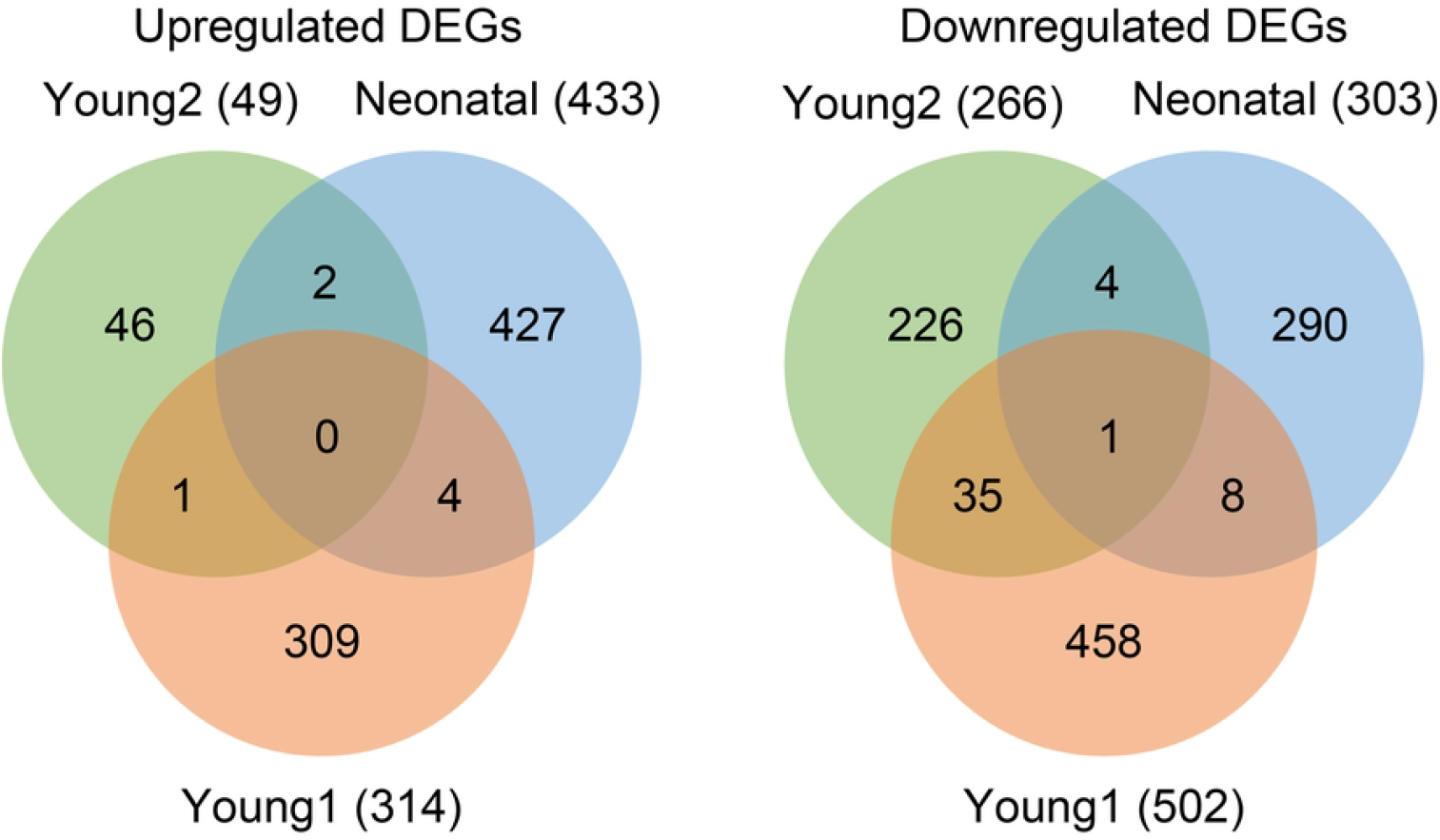
Venn diagrams of differentially expressed gene dataset after sevoflurane exposure. Transcriptome array data of the hippocampus of young mice treated with sevoflurane were compared with data from two studies: one by Yamamoto et al. [10] (referred to as Young 1) and one by Song et al. [7] (referred to as Young 2). The green cycle represents the data from Young 2, the orange cycle represents the data from Young 1, and the blue cycle represents the data from our neonatal mouse hippocampus.

### Neonatal sevoflurane exposure induced cognitive deficits exhibited during the Morris water maze test

Because the transcriptomic profile was significantly altered after exposure to sevoflurane, we investigated how multiple exposures to sevoflurane during the neonatal period affected the behavior of young mice on the Morris water maze test. During the probe trial, mice previously exposed to sevoflurane spent significantly less time in the target area than the control group (Fig 5A). Additionally, the sevoflurane group had longer first-time target latency than the control group (Fig 5B). Although there was no significant difference in the frequency of crossing the target area between the sevoflurane and control groups, there was a tendency for reduced frequency in the sevoflurane group. During the training trial, mice exposed to sevoflurane required more time to find the platform compared to that for the control mice when they were trained on days 4 and 5 (Fig 5D). These results indicate that exposure to sevoflurane impaired the ability of mice to learn and memorize.

**Fig 5.**
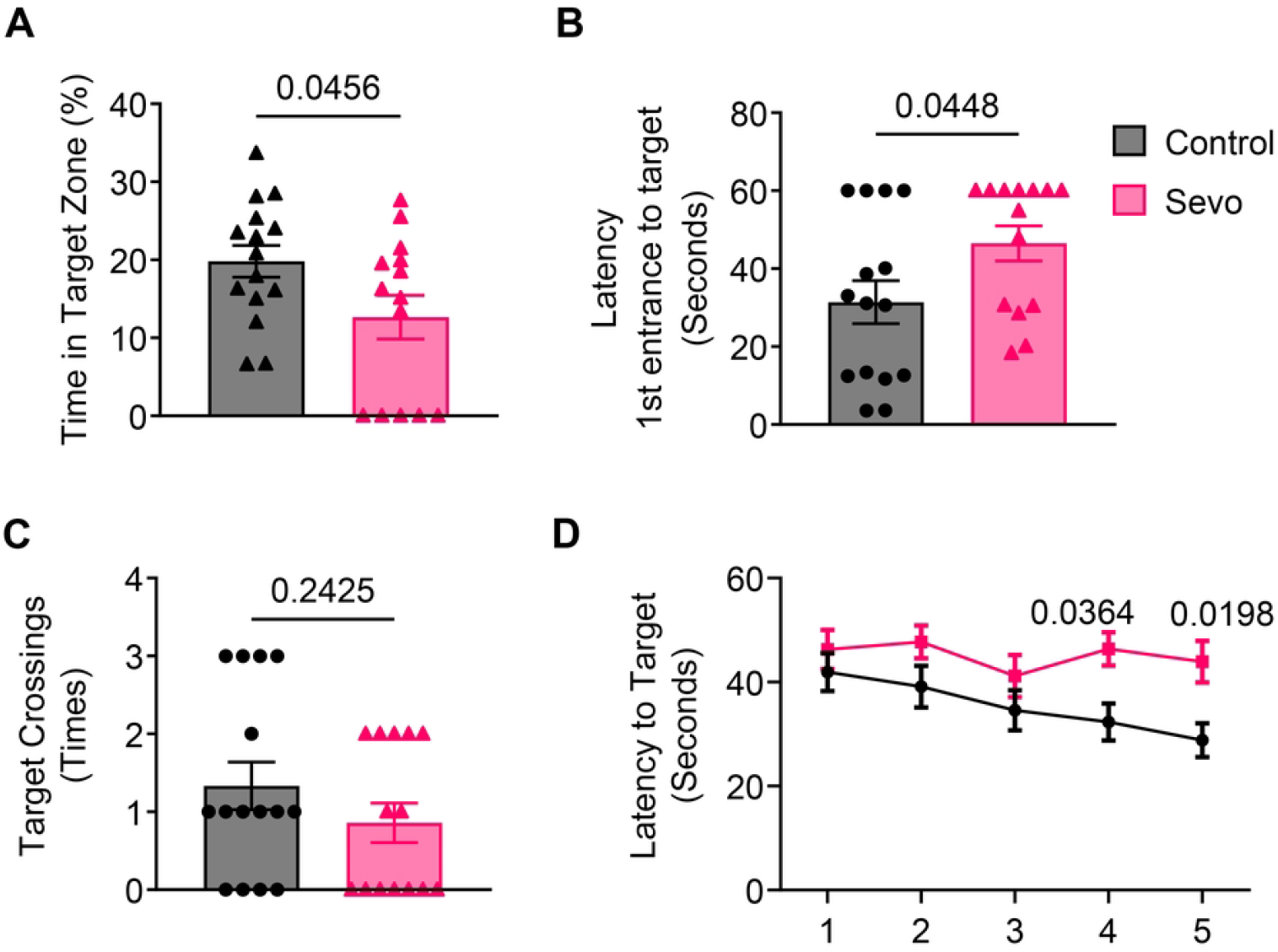
Results of Morris water maze test (days 1−6) and spatial exploration experiment (day 7, after removal of the platform). A) Proportion of time spent in the target area within 1 min. B) Time to first arrive at the platform. C) Number of times traversed across the platform zone. D) Time to arrival at the platform (escape latency) in the Morris water maze test.

### Neonatal sevoflurane exposure caused reduced synaptic protein expression levels

After discovering deficits in learning and memory after sevoflurane treatment, we attempted to elucidate the molecular mechanism underlying these occurrences. We analyzed the protein expression levels of synaptic-related genes. Our results showed that the expression levels of PSD95, SNAP25, and Bag3 were significantly decreased after sevoflurane exposure. The phosphorylation level of Bag3 increased after sevoflurane treatment (Fig 6A, B). In agreement with the Western blotting results, immunofluorescence staining confirmed that the expression of Bag3 in the hippocampus and cortex was significantly decreased with sevoflurane exposure (Fig 6C).

**Fig 6.**
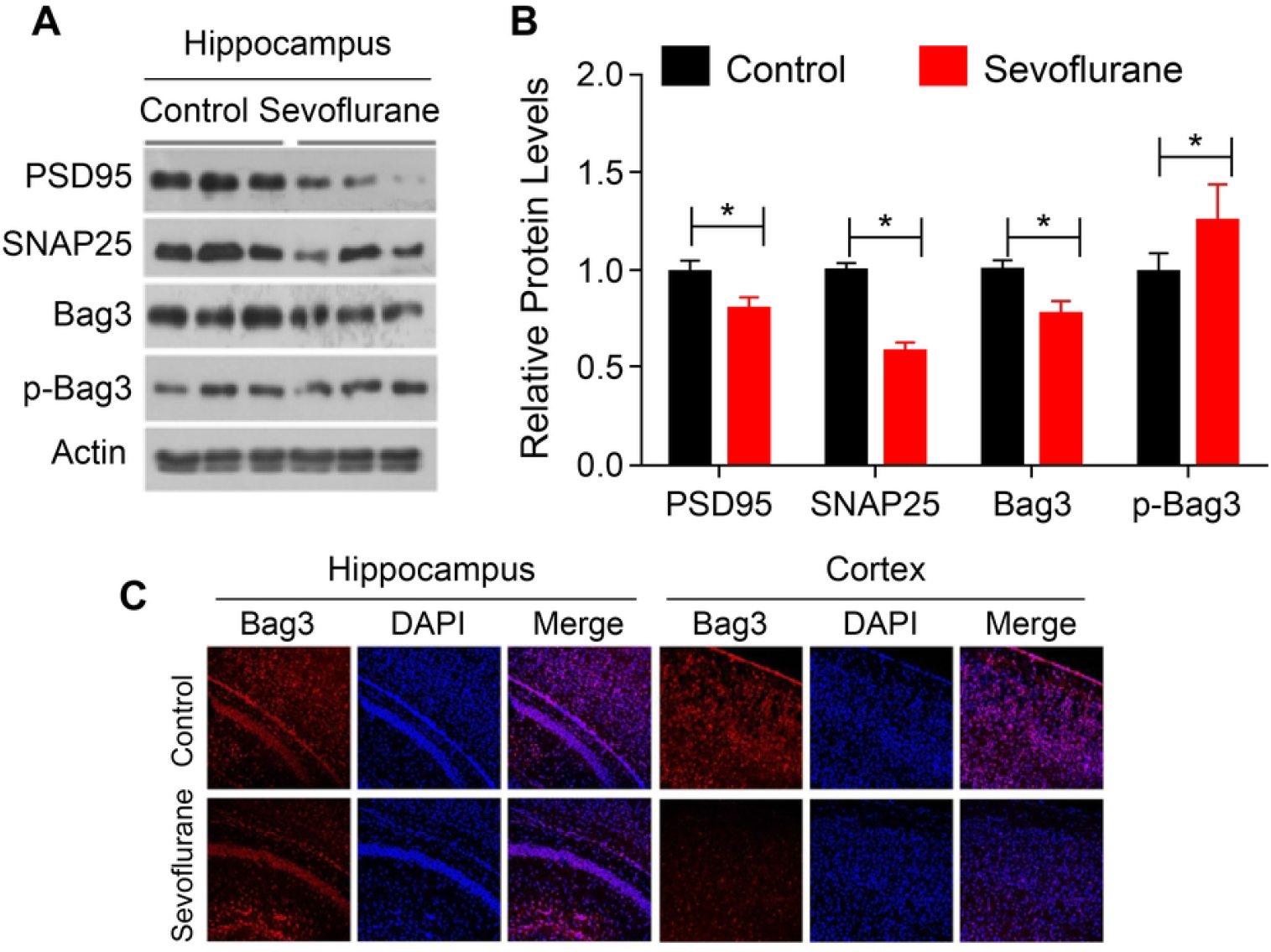
Western blotting and immunofluorescence staining. A, B). Western blotting reveals the effect of sevoflurane on the levels of synapse-associated proteins in the hippocampus. C) Immunofluorescence staining results. Bag3 expression decreased in the hippocampus and cerebral cortex after exposure to sevoflurane.

## Discussion

Sevoflurane has an important role in pediatric anesthesia; however, sevoflurane anesthesia can cause the development of cognitive dysfunction in children [11]. The precise molecular mechanism of sevoflurane exposure that affects brain development and cognitive dysfunction is not completely understood. Therefore, we performed RNA-seq of the neonatal hippocampus. This is the first study to assay the neonatal hippocampus transcriptome and investigate the molecular mechanism of memory deficits of neonatal mice exposed to sevoflurane. We identified 836 DEGs in the neonatal hippocampus after multiple exposures to sevoflurane compared to an unexposed control group. More genes were significantly upregulated (433) than downregulated (303). The most strongly upregulated genes were *ALB, AMBP, AHSG*, and *FGA*. In contrast, the most robustly downregulated genes were *TTR, TMEM72*, and *ISL1*. Significant changes in these genes indicate that sevoflurane exposure alters hippocampal development and is involved in axonal development. *TTR* binds to and distributes thyroid hormones in the blood and cerebrospinal fluid, which are critical to oligodendrocyte development. *AHSG* is highly expressed during brain development and is associated with brain development and immune function in the brain. The GO term enrichment analysis further indicated that these genes play a vital role in Parkinson’s disease-related gene expression induced by sevoflurane. Furthermore, based on the GO term analysis results, cellular components, biological processes, and molecular functions in the neonatal hippocampus were significantly altered. Moreover, the KEGG pathway analysis indicated the AMPK signaling pathway as one of the most important pathways involved in the downregulation of genes. Previous reports have revealed that the AMPK pathway is involved in sevoflurane exposure and Alzheimer’s disease mouse models. To further investigate the DEGs, we compared our dataset with two published datasets acquired as a result of studying the hippocampus in young mice after neonatal exposure to sevoflurane. Through this comparison, we found that the DEGs in the neonatal hippocampus were significantly different from those in the two published datasets. These data indicate that sevoflurane affects specific genes involved in hippocampal development.

Experimental evidence suggests that extensive neurodegeneration and cognitive dysfunction may occur in rodents and non-human primates exposed to sevoflurane [12]. The main mechanisms of sevoflurane-induced neurotoxicity discovered thus far include neuronal cell death, neuronal cell injury, neural circuit composition and plasticity, tau phosphorylation, and neuroendocrine effects [13]. Synaptic plasticity also has an essential role in the development of neural circuits, and impaired synaptic plasticity can lead to major neuropsychiatric disorders, especially hippocampal-dependent memory impairment [14]. The Morris water maze test has been widely used to investigate hippocampal-dependent cognitive functions in young mice. The present results showed that the escape latency and platform-seeking time were significantly prolonged, the residence time in the target area was shortened, and the number of crossings was reduced in mice with sevoflurane exposure. These results demonstrated that sevoflurane exposure in neonatal mice causes memory deficits in young mice.

We evaluated synapse-related protein levels in the hippocampus after sevoflurane modeling and found that SNAP25 protein levels were significantly decreased after sevoflurane exposure. Previous studies have demonstrated that SNAP25 is involved in synaptic vesicle docking, fusion, recycling, and neurotransmitter exocytosis, suggesting that sevoflurane may interfere with normal synaptic signaling [15]. PSD95 is another synaptic-related candidate that was significantly reduced after sevoflurane exposure. Consistent with previous findings, sevoflurane reduced the expression of PSD95 in the hippocampus of neonatal rats [16–18]. Therefore, sevoflurane may lead to synaptic plasticity dysfunction and induce abnormal neural circuits. Additionally, we estimated the total and phosphorylated Bag3 levels after sevoflurane exposure. Bag3 is an emerging autophagy regulator that interacts with heat shock protein 70 (HSP70) to maintain protein homeostasis [19], which is localized to neurites and synaptic neurons in the prefrontal cortex and hippocampus. Recently, several studies have suggested that Bag3 plays a vital role in synaptic function by regulating synaptic protein degradation [20]. Furthermore, sevoflurane can abnormally activate atypical cyclin-dependent kinase 5 (CDK5) [17]. CDK5-mediated phosphorylation of residue S297 promotes the degradation of Bag3, resulting in HSP70-mediated aberrant degradation of synaptic proteins and neurotransmitter receptors, thereby impairing synaptic plasticity and long-term potentiation in neurons. We found that the expression of Bag3 was decreased and that of p-Bag3 was increased in the sevoflurane group compared to that in the control group, suggesting that CDK5 phosphorylated and promoted Bag3 degradation, which may be involved in sevoflurane-induced nervous system dysfunction.

## Conclusion

The preset genome-wide transcriptional analysis of the neonatal mouse hippocampus demonstrated that sevoflurane exposure significantly changed the gene expression in the neonatal hippocampus. Moreover, the Morris water maze test results suggested that neonatal sevoflurane anesthesia persistently impair learning and memory during adolescence. The biochemical analysis showed that reductions in PSD95, SNAP25, and Bag3 protein levels and increased phosphorylation of Bag3 under sevoflurane exposure may induce synaptic dysfunction, thereby causing memory impairment in mice. This study revealed a transcriptome-wide profile in the neonatal mice hippocampus by performing a biochemical analysis and behavioral testing after multiple exposures to sevoflurane during the neonatal period, thus providing a better understanding and potential explanation of the mechanisms of the impact of anesthesia on the brain function in young children exposed to sevoflurane.

## Acknowledgments

This work was supported by the First Affiliated Hospital of Xiamen University, Department of Anesthesiology, which funded the work in all initial stages. Equipment and materials were provided by the Fujian Provincial Key Laboratory of Neurodegenerative Disease and Aging Research, Institute of Neuroscience. We would like to thank Editage (www.editage.cn) for English language editing.

